## Supplemental data file for "Efficient pheromone navigation via antagonistic detectors in *Caenorhabditis elegans* male"

**Supplementary Table 1**

| REAGENT or RESOURCE | SOURCE | IDENTIFIER |
|---|---|---|
| **Bacterial and virus strains** | | |
| *E. coli* | *Caenorhabditis* Genetics Center (CGC) | WormBase: OP50; WormBase: WBStrain00041969 |
| **Experimental models: Organisms/strains** | | |
| *him-5 (e1490) V* | CGC | CB4088 (male enriched mutant) |
| *srd-1(eh1)* | CGC | CB5414 (*srd-1* mutant) |
| *syEx1972 [srd-1P::mCherry]; syEx1974 [unc-119::gfp]* | This study | PS10451 (*srd-1* reporter; pan-neural marker) |
| *syEx1973 [rab-3P::mCherry]; syIs912[srd-1P::Gal4(sk)::VP64]; syIs300 [15xUAS::GFP]* | This study | PS10448 (*srd-1* driver; GFP effector; pan-neural marker) |
| *syEx1972 [srd-1P::mCherry]; syIs913 [pkd-2P::Gal4(sk)::VP64]; syIs300 [15xUAS::GFP]* | This study | PS10449 (*srd-1* reporter; *pkd-2* driver; GFP effector) |
| *syIs912[srd-1P::Gal4(sk)::VP64]* | This study | PS9477 (*srd-1* driver) |
| *syIs914[srd-1P::Gal4(sk)::VP64]* | This study | PS9478 (*srd-1* driver) |
| *syIs912[srd-1P::Gal4(sk)::VP64]; syIs300[UAS::GFP]* | This study | PS9473 (*srd-1* driver; GFP effector) |
| *syIs913[pkd-2P::Gal4(sk)::VP64]; syIs300 [15xUAS::GFP]* | This study | PS9681 (*pkd-2* driver; GFP effector) |

| | | |
|---|---|---|
| *syIs838[lin-48dP::NLS::cGAL (DBD)::gp41-1::N-intein::let-858 3'UTR, srd-1P::NLS::gp41-1::C-intein::cGAL(AD)::let-858 3'UTR]* | This study | PS10007 (PHD driver) |
| *syIs838[lin-48dP::NLS::cGAL (DBD)::gp41-1::N-intein::let-858 3'UTR, srd-1P::NLS::gp41-1::C-intein::cGAL(AD)::let-858 3'UTR]; syIs300[15xUAS::GFP]* | This study | PS9573 (PHD driver; GFP effector) |
| *syIs888 [gpa-14P::NLS::cGAL (DBD)::gp41-1::N-intein::let-858 3'UTR, rig-3P::NLS::gp41-1::C-intein::cGAL(AD)::let-858 3'UTR]* | This study | PS10179 (AVA driver) |
| *syIs888[gpa-14P::NLS::cGAL (DBD)::gp41-1::N-intein::let-858 3'UTR, rig-3P::NLS::gp41-1::C-intein::cGAL(AD)::let-858 3'UTR] ]; syIs300[15xUAS::GFP]* | This study | PS10087 (AVA driver; GFP effector) |
| *syIs564 [odr-10P::Gal4(sk)::VP64]; syIs300 [15xUAS::GFP]* | This study | PS8293 (AWA driver; GFP effector) |
| *syIs340 [15xUAS::hChR (H134R)::EYFP::let-858 3'UTR]; him-5 (e1490) V* | This study | PS9782 (Chrimson effector; *him-5*) |
| *syIs340 [15xUAS::hChR (H134R)::EYFP::let-858 3'UTR]; him-5 (e1490) V* | This study | PS7043 (Chrimson effector) |
| *syIs371[15xUAS::HisCL::SL2::GFP::let-858 3'UTR]* | This study | PS7199 (HisCL effector) |
| *syIs612[15xUAS::GCaMP7b::SL2::mKate2]* | This study | PS9046 (GCaMP7b effector) |
| *syIs912[srd-1P::Gal4(sk)::VP64]; syIs612[15xUAS::GCaMP7b::SL2::mKate2]* | This study | PS9680 (*srd-1* driver; GCaMP7b effector) |
| *syIs912[srd-1P::Gal4(sk)::VP64]; syIs371[15xUAS::HisCL::SL2::GFP::let-858 3'UTR]* | This study | PS10005 (*srd-1* driver; HisCl effector) |
| *syIs912[srd-1P::Gal4(sk)::VP64]; syIs340 [15xUAS::hChR (H134R)::EYFP::let-858 3'UTR]; him-5 (e1490) V* | This study | PS9990 (*srd-1* driver; Chrimson effector) |
| *syIs838[lin-48dP::NLS::cGAL(DBD)::gp41-1::N-intein::let-858 3'UTR, srd-1P::NLS::gp41-1::C-intein::cGAL(AD)::let-858 3'UTR]; syIs371[15xUAS::HisCL::SL2::GFP::let-858 3'UTR]* | This study | PS10189 (PHD driver; HisCL effector) |
| *syIs838[lin-48dP::NLS::cGAL(DBD)::gp41-1::N-intein::let-858 3'UTR, srd-1P::NLS::gp41-1::C-intein::cGAL(AD)::let-858 3'UTR]; syIs340 [15xUAS::hChR(H134R)::EYFP::let-858 3'UTR]; him-5(e1490)V.* | This study | PS10187 (PHD driver; Chrimson effector) |
| *syIs888 [gpa-14P::NLS::cGAL (DBD)::gp41-1::N-intein::let-858 3'UTR, rig-3P::NLS::gp41-1::C-intein::cGAL(AD)::let-858 3'UTR]; syIs612[15xUAS::GCaMP7b::SL2::mKate2]* | This study | PS10181 (AVA driver; GCaMP7b effector) |
| *syIs564 [odr-10P::Gal4(sk)::VP64]; syIs340 [15xUAS::hChR (H134R)::EYFP::let-858 3'UTR]* | This study | PS9387 (AWA driver; Chrimson effector) |
| *hpIs675 [rgef-1P::GCaMP6s::3xNLS::mNeptune + lin-15(+)]; lin-15(n765) X; him-5 (e1490) V* | This study | ZM9627 (pan-neuronal nucleus localized mNeptune and GCaMP6s) |
| *hpIs675 [rgef-1P::GCaMP6s::3xNLS::mNeptune + lin-15(+)]; lin-15(n765) X; him-5 (e1490) V; srd-1(eh1)* | This study | PS10450 (pan-neuronal nucleus localized |

| | | |
|---|---|---|
| | | mNeptune and GCaMP6s in *srd-1(eh1)* background) |
| *lite-1(ce314); JAC66[ ift-20P:BFP; eat-4P::cyOFP1; unc-17P::mScarlet: acr-5P::BFP; hpIs675 [rgef-1P::GCaMP6s::3xNLS::mNeptune + lin-15(+)]; lin-15(n765) X* | This study | ADS1046 (pan-neuronal nucleus localized mNeptune, GCaMP6s, and neuron ID markers) |
| *lite-1(ce314); JAC66[ ift-20P:BFP; eat-4P::cyOFP1; unc-17P::mScarlet: acr-5P::BFP; hpIs675 [rgef-1P::GCaMP6s::3xNLS::mNeptune + lin-15(+)]; lin-15(n765) X; syIs912[srd-1P::Gal4(sk)::VP64]; syIs340 [15xUAS::hChR(H134R)::EYFP::let-858 3'UTR]; him-5(e1490).* | This study | PS10153 (pan-neuronal nucleus localized mNeptune, GCaMP6s, neuron ID markers; *srd-1* driver; Chrimson effector) |
| *Kp1368 [myo-2P::NLS::mCherry]; syIs371[15xUAS::HisCL::SL2::GFP::let-858 3'UTR]* | This study | PS8720 (muscle driver; HisCl effector) |
| *syIs334 [rabP::GAL4::gp41::N::let-858 3' UTR]; syIs340 [15xUAS::hChR(H134R)::EYFP::let-858 3'UTR]; him-5(e1490).* | This study | PS8026 (pan-neural driver; Chrimson effector) |
| **Software and algorithms** | | |
| Fiji (Fiji Is Just ImageJ) — v2.16.0 | Schindelin et al., 2012 [117] | https://imagej.net/software/fiji/ |
| MaMuT 0.27 | Wolff et al., 2018 [118] | https://imagej.net/plugins/mamut/ |
| Target track (2018) | Park et al., 2018 [119] | https://www.nature.com/articles/s41592-023-02096-3 |
| Ilastik software v1.4.0.post1 | Berg et al., 2019 [120] | https://www.ilastik.org |
| **Other** | | |
| cGAL-UAS bipartite expression toolkit | Nava et al., 2023 [121] | https://www.pnas.org/doi/full/10.1073/pnas.2221680120 |

22 **Supplementary Figure and Legends**

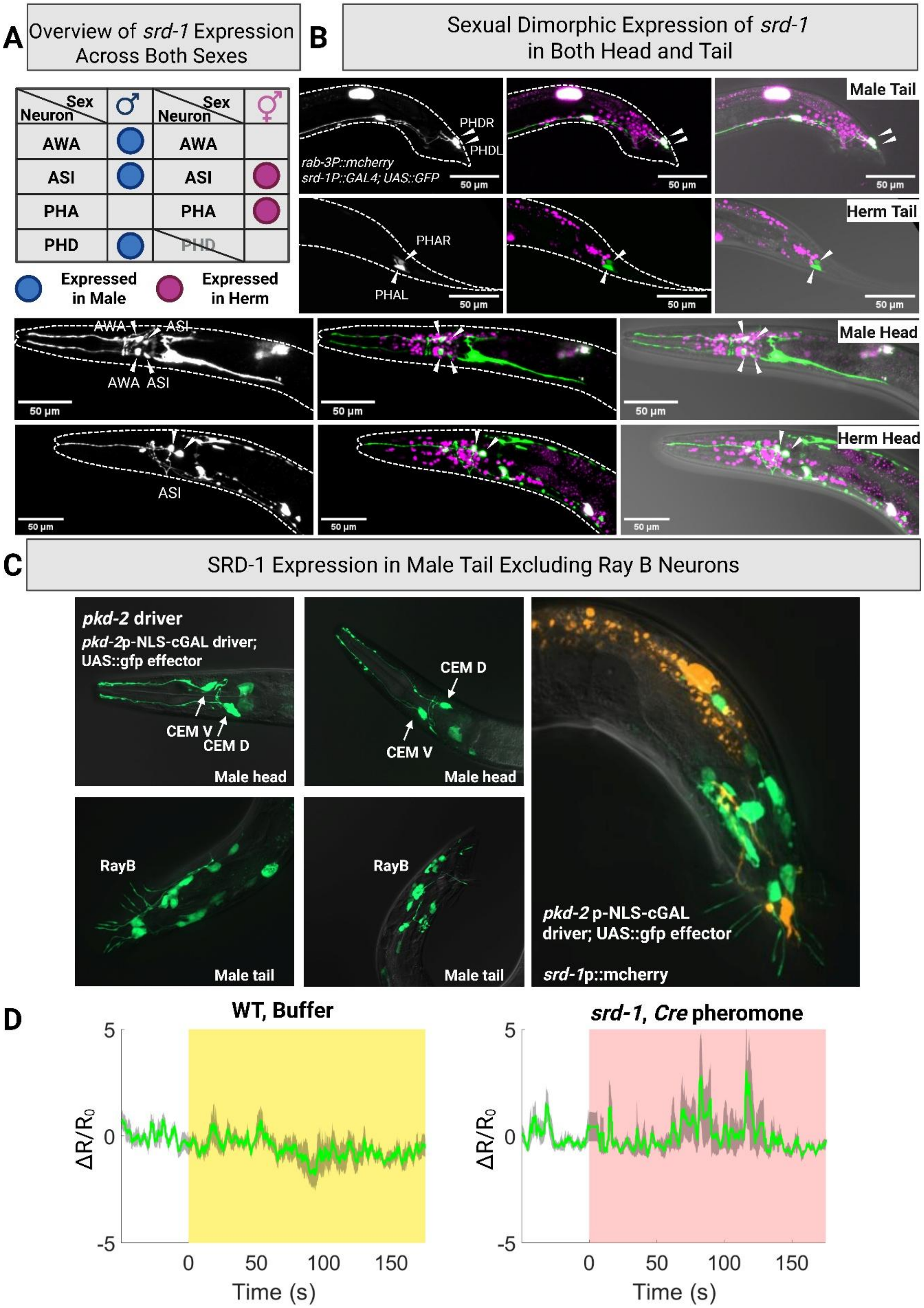

**Figure S1: Localization of SRD-1 expression in the male tail, excluding ray B neurons. (A)** Overview of *srd-1* expression patterns in both sexes. **(B)** Sexual dimorphic expression of *srd-1* in both head and tail: *srd-1* GFP reporter expression in AWAs, ASIs and male-specific neuron PHDs in males and in ASIs and PHAs in hermaphrodites. *rab-3P::mCherry* marks all neurons; an integrated cGAL-UAS line *(srd-1P*::GAL4; UAS::GFP*)* reports *srd-1* expression. **(C)** Visualization of *pkd-2* expression in the male tail region using a cGAL-UAS system: *pkd-2P*-NLS-cGAL driver combined with a UAS::gfp effector. PKD-2 is known to be expressed in CEM neurons in the head and Ray B neurons in the tail. The figure presents two representative males. Absence of overlap between PKD-2 (GFP) and SRD-1 (mCherry) expressions in the male tail region, utilizing *pkd-2p*-NLS-cGAL driver and UAS::*gfp* effector in combination with *srd-1P*::mCherry, demonstrating distinct expression domains. **(D)** Zoomed view of Figure 1C first and last panel ( y-axis -5 to 5). Pan-neuronal calcium imaging of male PHD neurons expressing *rgef-1P*::GCaMP6s, with *rgef-1P*::mNeptune marking all neuronal somata. WT males received M9 buffer (control); *srd-1* mutant males received volatile sex pheromone. Traces show mean $\Delta R/R_0$ (R = GCaMP/mNeptune fluorescence ratio) ± s.e.m.; n = 5–7 animals per condition. The green trace denotes the mean calcium ratio ($\Delta R/R_0$). Shading denotes pre-stimulus (white), M9 control buffer application (yellow), and pheromone application (pink). Source data are provided as a Source Data file.

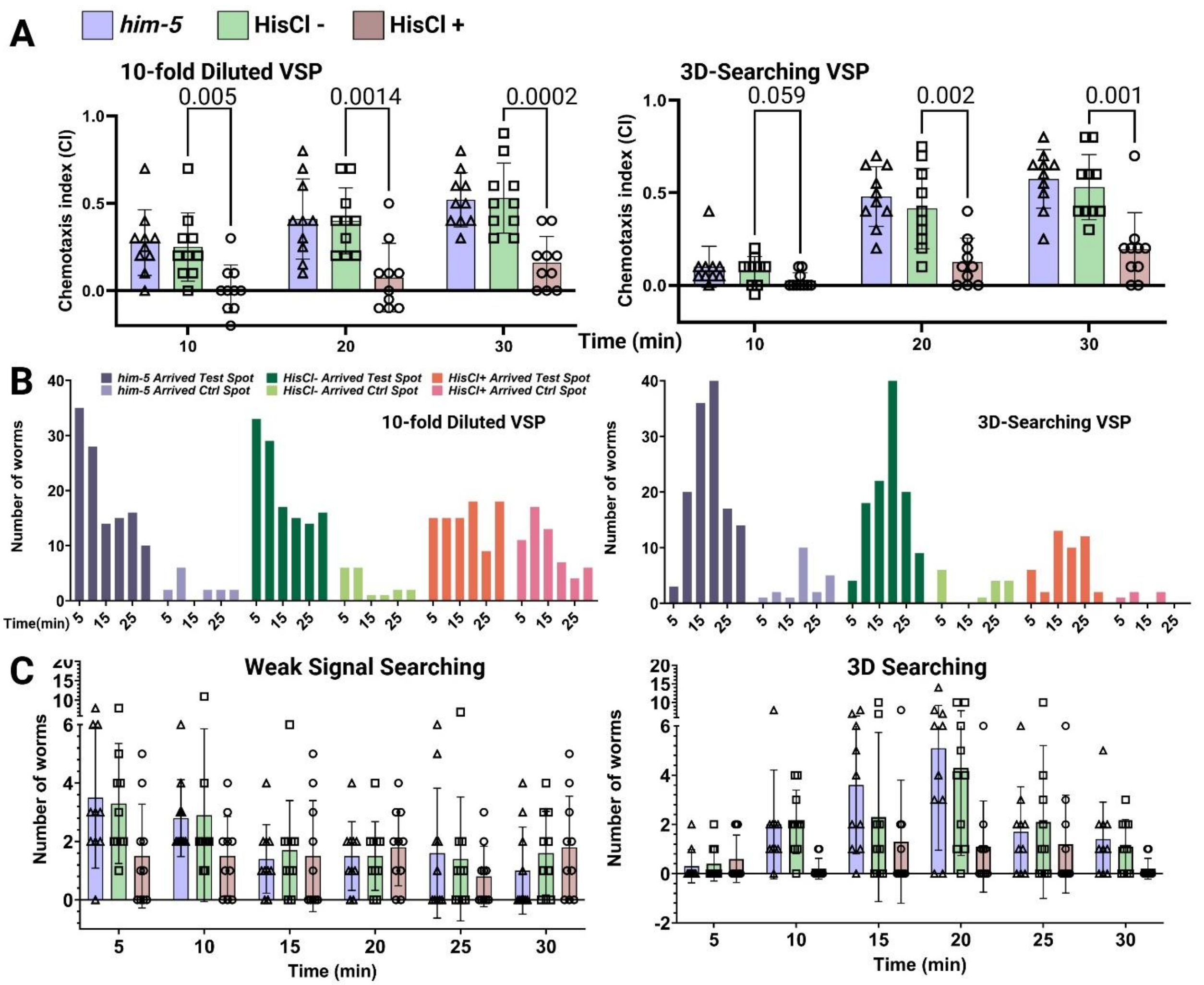


**Figure S2: Chemotaxis performance of *him-5* males in weak signal and 3D searching. (A)** Temporal profile of the chemotaxis index (CI). The distribution of worms at various intervals following stimulus shows a consistently low chemotaxis index in PHD-inhibited males across different time point. **(B)** Worm counts at each spot across time points. Different navigational deficits in PHD-inhibited males across different exploration tasks: in weak signal tasks, their even distribution between test and control spots led to a low chemotaxis index; in 3D tasks, the low index resulted from fewer males reaching the target. Inhibition was achieved by crossing the indicated PHD neurons-specific driver line to 15×UAS::HisCl::SL2::GFP and assaying on 10 mM histamine; *him-5* and genotype-matched animals maintained on histamine-free plates served as controls (See Methods). **(C)** Presents the number of worms reaching a pheromone target spot at various time points during weak signal searching and 3D searching assays. The analysis of worm distribution at multiple time points after the introduction of the stimulus indicated a consistently low number of the PHD-inhibited males arrived at the test spot across all intervals, highlighting their compromised chemotactic ability. **The sample size for each assay consisted of 200 worms, 20 worms per trials, 10 trails.** Two-tailed unpaired t-test. Error bars represent the s.e.m. Exact P values are indicated on the figure. Source data are provided as a Source Data file.

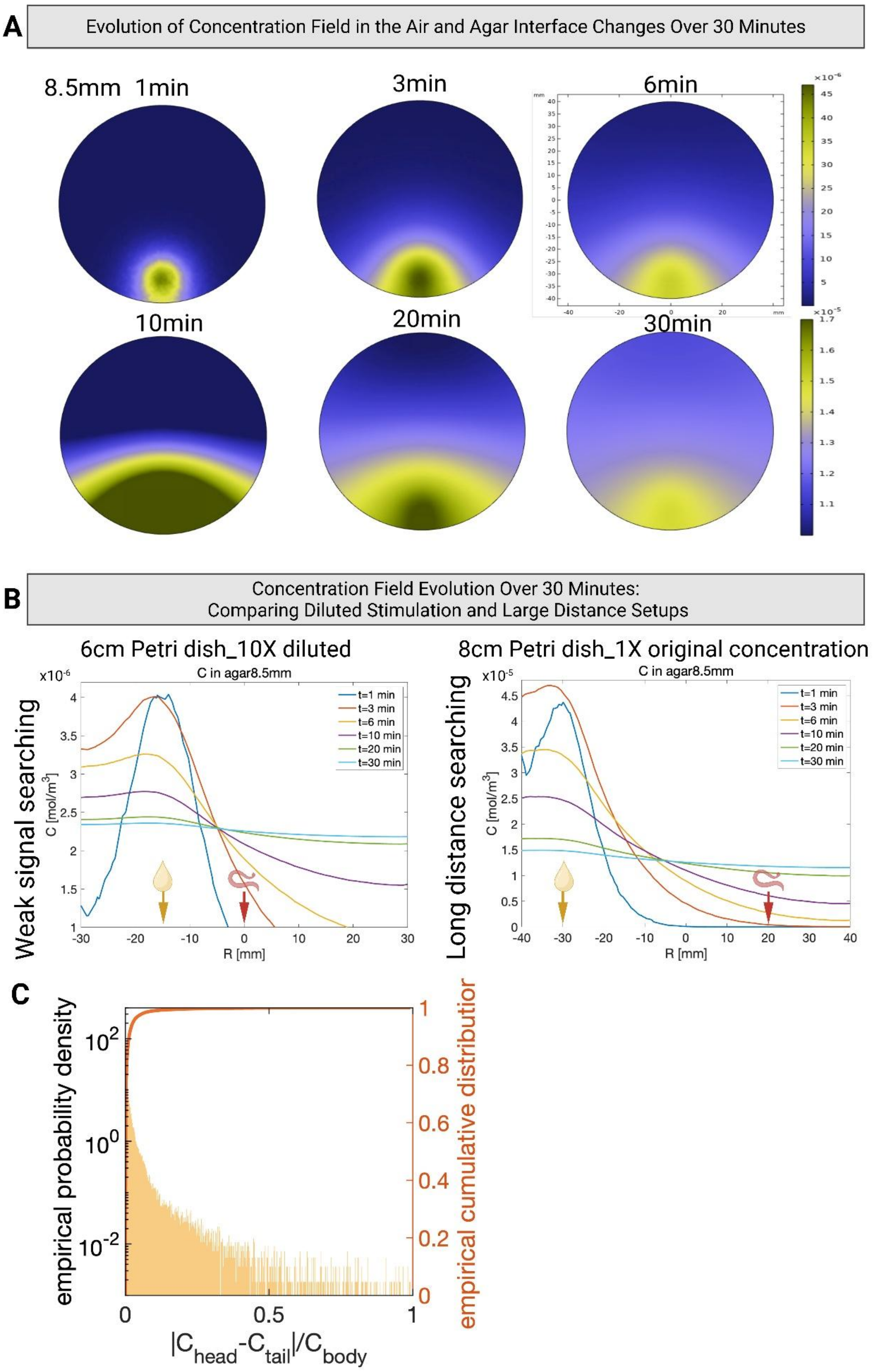
A
Evolution of Concentration Field in the Air and Agar Interface Changes Over 30 Minutes
8.5mm 1min
3min
6min
10min
20min
30min
B
Concentration Field Evolution Over 30 Minutes:
Comparing Diluted Stimulation and Large Distance Setups
6cm Petri dish_10X diluted
8cm Petri dish_1X original concentration
Weak signal searching
Long distance searching
C in agar8.5mm
C [mol/m^3]
R [mm]
t=1 min
t=3 min
t=6 min
t=10 min
t=20 min
t=30 min
C
empirical probability density
empirical cumulative distributior
|Chead-Ctail|/Cbody

73

**Figure S3: Visualization and dynamics of pheromone concentration fields in chemoattraction assays. (A)** Captures the evolution of the concentration field at the air and agar interface (8.9mm) over a 30-minute duration. The color scales for 1min, 3min, and 6min intervals are consistent, as are those for 10min, 20min, and 30min intervals, as indicated on the right side of the panel, illustrating the dynamic changes in pheromone distribution. **(B)** Compares the evolution of the concentration field over 30 mins between diluted stimulation and large distance setups. The concentration field simulation shows that in scenarios where males are positioned far from the pheromone source, they experience areas of low absolute concentration lacking a clear gradient. In contrast, proximity to a diluted pheromone source results in a Gaussian distribution with a discernible gradient, despite the overall low concentration level, indicating different challenges in pheromone detection and navigation based on distance and concentration. **(C)** Distribution of concentration difference between head and tail sensory neurons, normalized by the concentration of body center and averaged over all spatial locations on the plate, the entire simulation time, and worm orientation. Yellow bars correspond to the left y-axis, showing the empirical probability density distribution (EPDF). We show the y-axis in log-scale to assist visualization since the EPDF is strongly concentrated near 0. The orange line corresponds to the right y-axis, showing the empirical cumulative distribution function (ECDF). The 99% quantile is 0.1056. Created in BioRender. Wan, X. (2026) https://BioRender.com/cpobd04. Source data are provided as a Source Data file.

91

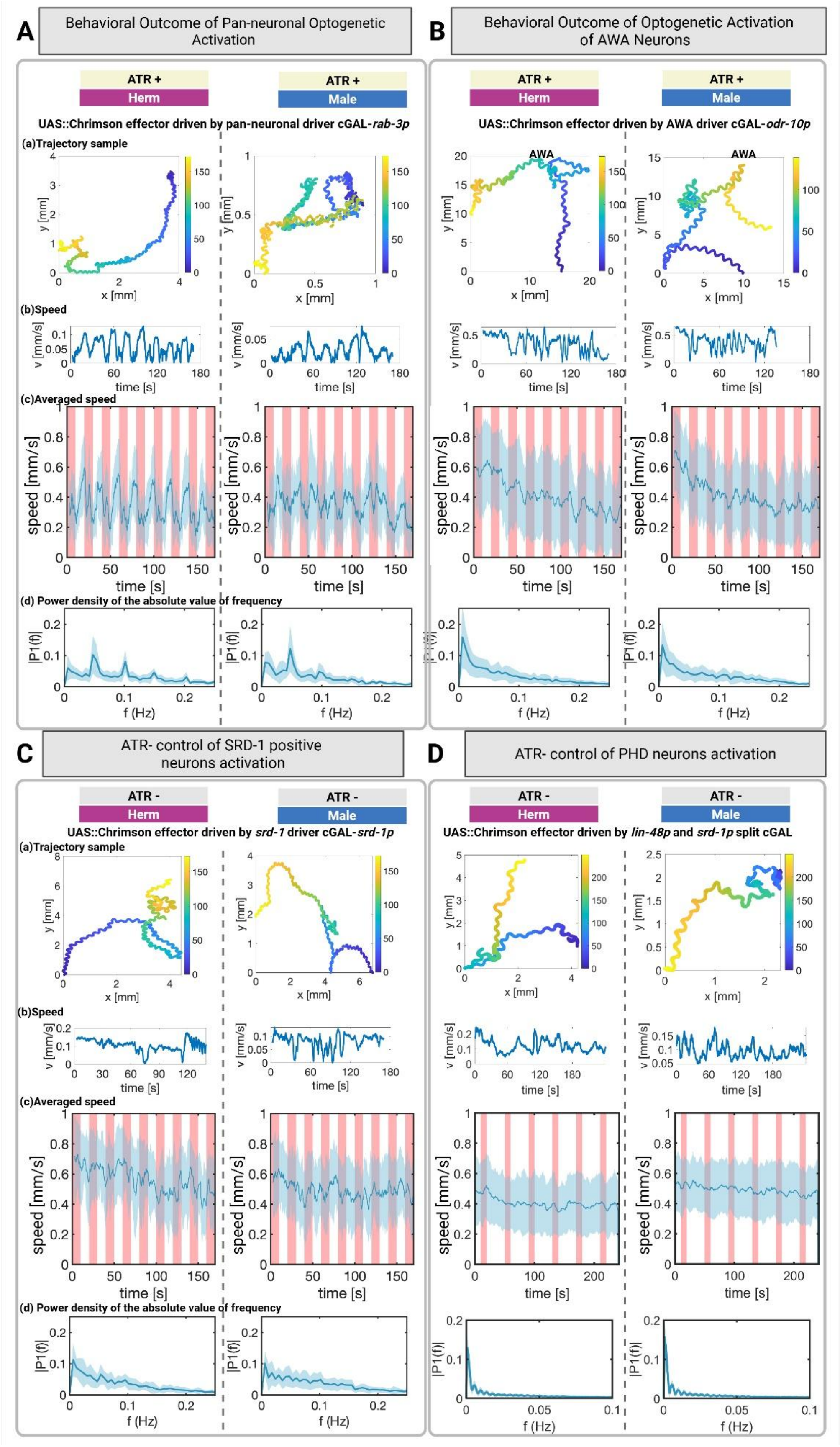

A
Behavioral Outcome of Pan-neuronal Optogenetic Activation
ATR +
Herm
ATR +
Male
UAS::Chrimson effector driven by pan-neuronal driver cGAL-*rab-3p*
(a)Trajectory sample
(b)Speed
(c)Averaged speed
(d) Power density of the absolute value of frequency
B
Behavioral Outcome of Optogenetic Activation of AWA Neurons
ATR +
Herm
ATR +
Male
UAS::Chrimson effector driven by AWA driver cGAL-*odr-10p*
AWA
C
ATR- control of SRD-1 positive neurons activation
ATR -
Herm
ATR -
Male
UAS::Chrimson effector driven by *srd-1* driver cGAL-*srd-1p*
(a)Trajectory sample
(b)Speed
(c)Averaged speed
(d) Power density of the absolute value of frequency
D
ATR- control of PHD neurons activation
ATR -
Herm
ATR -
Male
UAS::Chrimson effector driven by *lin-48p* and *srd-1p* split cGAL
x [mm]
y [mm]
v [mm/s]
time [s]
speed [mm/s]
|P1(f)|
f (Hz)

**Figure S4: Effects of Pan-Neural and AWA Neuron Activation on *C. elegans* Locomotion and ATR- Control of SRD-1 Positive Neuron and PHD Neuron Activation. (A)** Demonstrates global neuronal activation via the Chrimson system in *C. elegans*, employing a pan-neuronal promoter (*rab-3p*-NLS-cGAL driver) to drive UAS::Chrimson effector expression. This approach achieved effective neuronal stimulation across the nervous system in both male and female worms, leading to a temporary cessation of movement during light exposure, with immediate resumption of normal behavior upon cessation of the stimulus. The frequency of movement cessation closely followed the stimulation protocol, showing a dominant frequency at 0.05Hz, indicating the system's effectiveness in modulating worm locomotion through pan-neural activation. **(B)** AWA activation was achieved by expressing Chrimson in AWA using an *odr-10P* cGAL driver line crossed to a 15×UAS::Chrimson; animals were raised ± ATR and AWA neurons are optogenetically activated by red light during the assay. The figure includes (a)(b) a representative trajectory sample alongside its total velocity time series V(t), with the red line marking periods of light stimulation; (c) the averaged V(t) across 30 samples, highlighted by a red shade during light stimulation; and (d) the Fourier analysis of V(t), illustrating the differential response patterns to pan-neural versus AWA-specific activation. The sample size for the pan-neural activation assay included 50 worms per sex, while the AWA activation assay involved 110-120 worms per sex. Shaded regions indicate ± s.d.; solid lines are means. **(C)** Locomotor responses to SRD-1 positive neuron activation in both sexes of *C. elegans*. SRD-1 positive neurons (*srd-1P*-NLS-cGAL driver) were activated in both sexes using Chrimson (UAS::Chrimson effector). The control, without all-trans retinal (ATR), had no change in behavior, eliminating any light effects. **(D)** Locomotor responses to PHD neuron activation in both sexes of *C. elegans*. PHD neurons (split cGAL-UAS line: *lin-48dP*-NLS-cGAL (DBD); *srd-1P*-NLS-cGAL (AD) driver) were activated in both sexes using Chrimson (UAS::Chrimson effector). The control, without all-trans retinal (ATR), also had no change in behavior, eliminating any light effects. **(C) (D)** The sample size for the SRD-1 positive neuron and PHD neuron activation assay included 120 worms per sex (See ATR+ control in Figure 3). Source data are provided as a Source Data file.

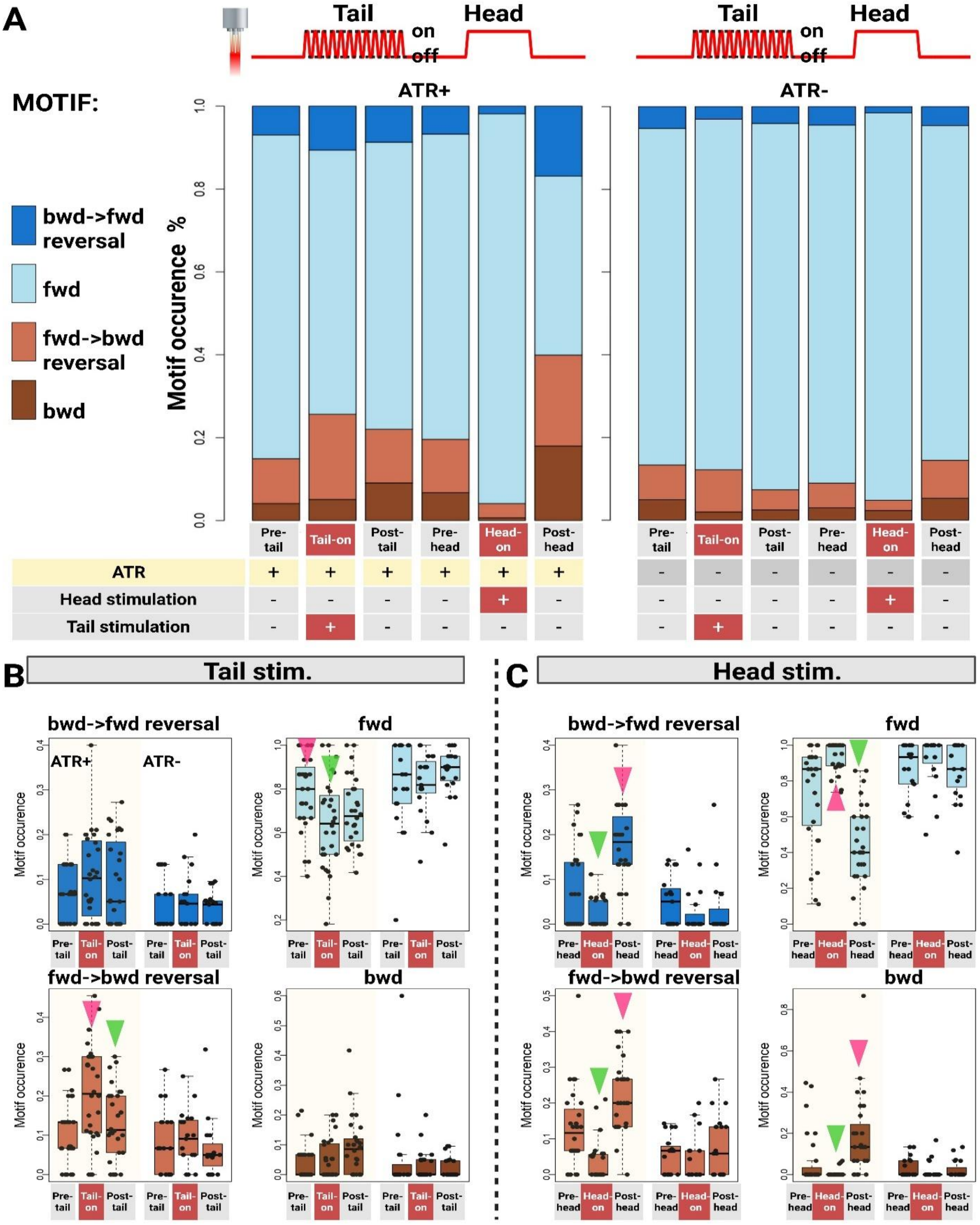
A
Tail
Head
on
off
ATR+
ATR-
MOTIF:
bwd->fwd reversal
fwd
fwd->bwd reversal
bwd
Motif occurence %
Pre-tail
Tail-on
Post-tail
Pre-head
Head-on
Post-head
ATR
Head stimulation
Tail stimulation
B
Tail stim.
C
Head stim.
Motif occurence

126

**Figure S5: Four-Motif Analysis: Differential locomotion responses to optogenetic stimulation of SRD-1 positive neurons in male *C. elegans.*** Utilizes the *srd-1* Chrimson cGAL-UAS system for precise optogenetic stimulation of SRD-1 positive neurons in the head and tail regions of male *C. elegans*, revealing distinct locomotive responses. Tail-targeted light pulses induced transitions from forward to backward movement and hesitancy in movement direction. Continuous head stimulation resulted in sustained forward motion and exploratory post-stimulation behaviors. Experiments included ATR+ (n=18) and ATR- (n=12) groups, with sequential recording periods before, during, and after stimulation in both regions, to assess the nuanced impact on locomotion. **(A)** Tail activation effects on locomotion. A short light pulse targeting the tail induced a notable transition from forward to backward movement and increased hesitancy in movement direction during stimulation. **(B)** Head activation and subsequent behaviors. Continuous head stimulation for 30 seconds prompted sustained forward movement during light exposure and immediate directional adjustments post-stimulation. After head stimulation, males engaged in self-exploratory behaviors, using their tails for body probing. Each worm was recorded for 30 seconds before tail stimulation, followed by a 2-second alternating light pulse targeting the tail over 12 cycles. Post-tail stimulation, there was another 30-second recording period before initiating head stimulation, then the head stimulation lasted for 30 seconds. The worm was then recorded for an additional 30 seconds after the light stimulation concluded. Error bars represent the s.e.m. Source data are provided as a Source Data file.

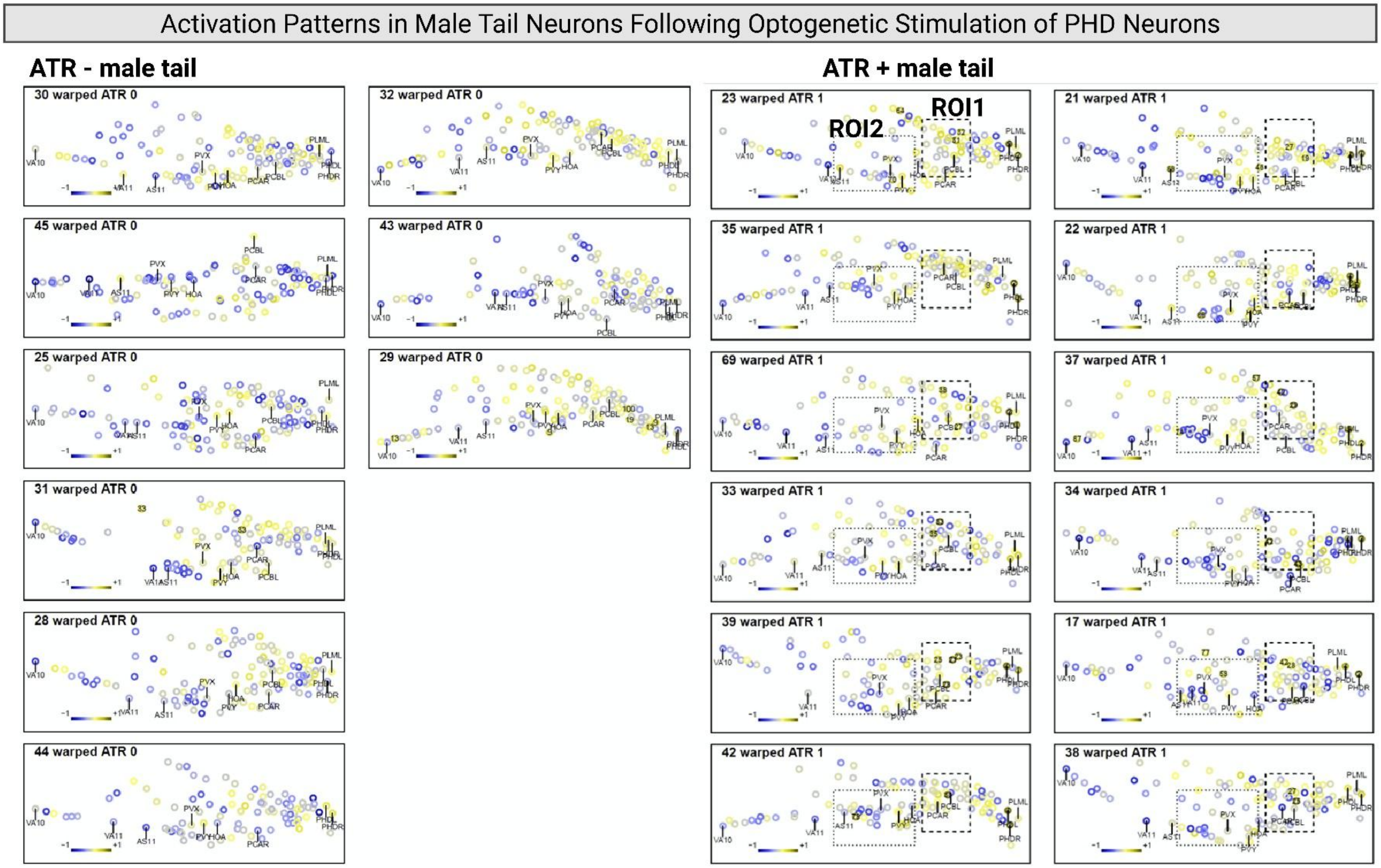


**Figure S6: Activation patterns observed in male tail neurons upon optogenetic stimulation of PHD neurons.** Visual analysis identified two regions, designated ROI1 and ROI2 in multiple males, which contain neurons with strong correlations to PHD activity in ATR+ group, but not ATR- group (correlation coefficient >0.55). Sample sizes: control (n=9) and experimental (n=12). Source data are provided as a Source Data file.

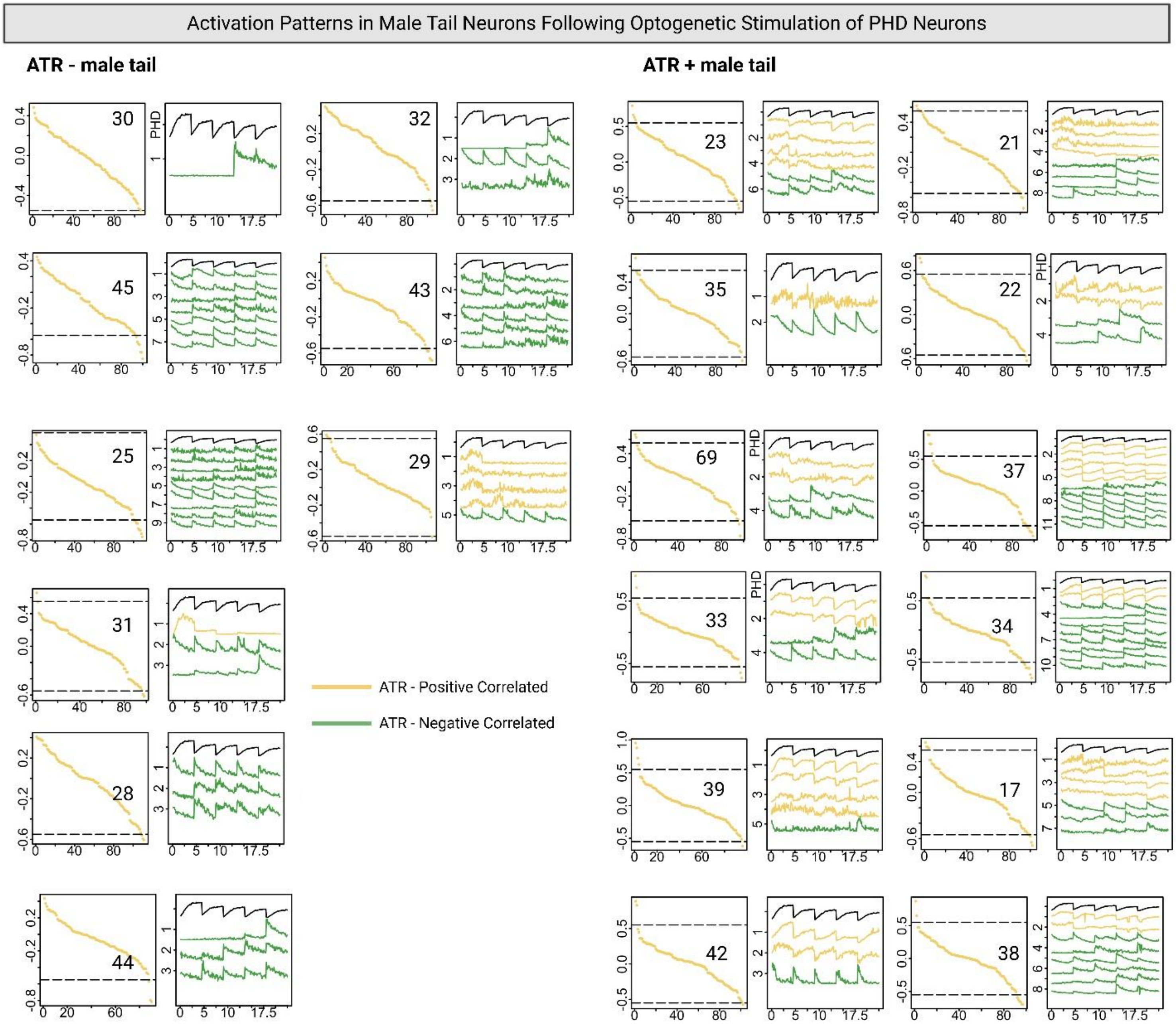


**Figure S7: Neural activity correlations with PHD optogenetic stimulation in tail regions.** Yellow and purple indicate positive correlations with target neuron activity in ATR- and ATR+ samples, respectively. Green denote negative correlations in ATR- and ATR+ samples, respectively. The dashed lines indicate the thresholds for positive and negative correlations. Sample sizes: control (n=9) and experimental (n=12). Source data are provided as a Source Data file.

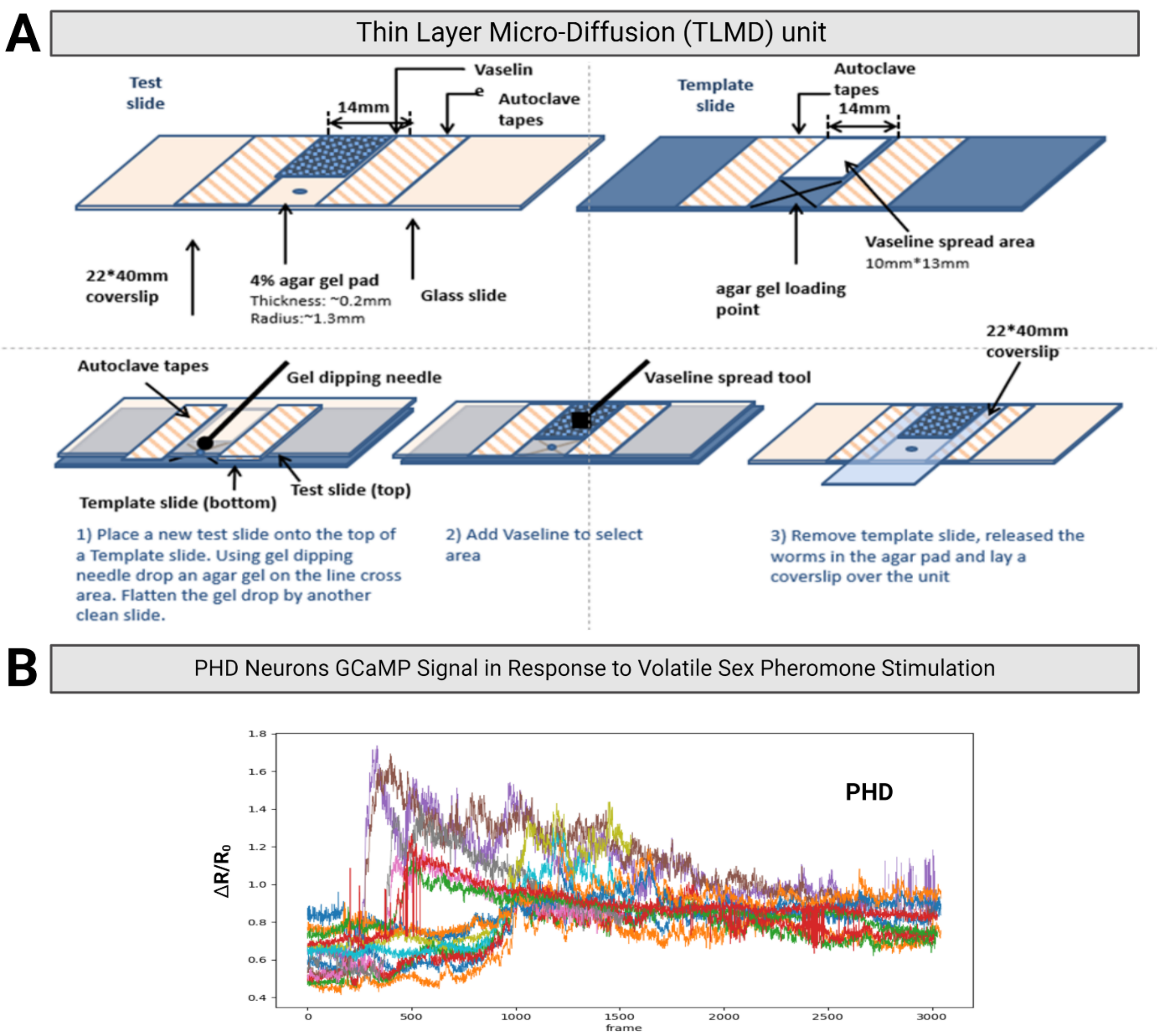


**Figure S8: TLMD unit for neuronal activation and calcium response measurement. (A)** Depicts the Thin Layer Micro-Diffusion (TLMD) unit setup for detecting changes in intracellular calcium signaling via GCaMP6s fluorescence intensity. The device consists of a 4% agarose pad placed between autoclave tapes, used with 0.2µl of 1mM levamisole to partially immobilize well-prepared transgenic worms. Test chemicals are introduced at 30 seconds, with rapid diffusion facilitated by capillary action. **(B)** Calcium responses of PHD neurons during 9.5 mins of pheromone stimulation, with individual animals distinguished by color coding. The sample sizes is 13 worms. Shaded areas represent the s.e.m. Source data are provided as a Source Data file.

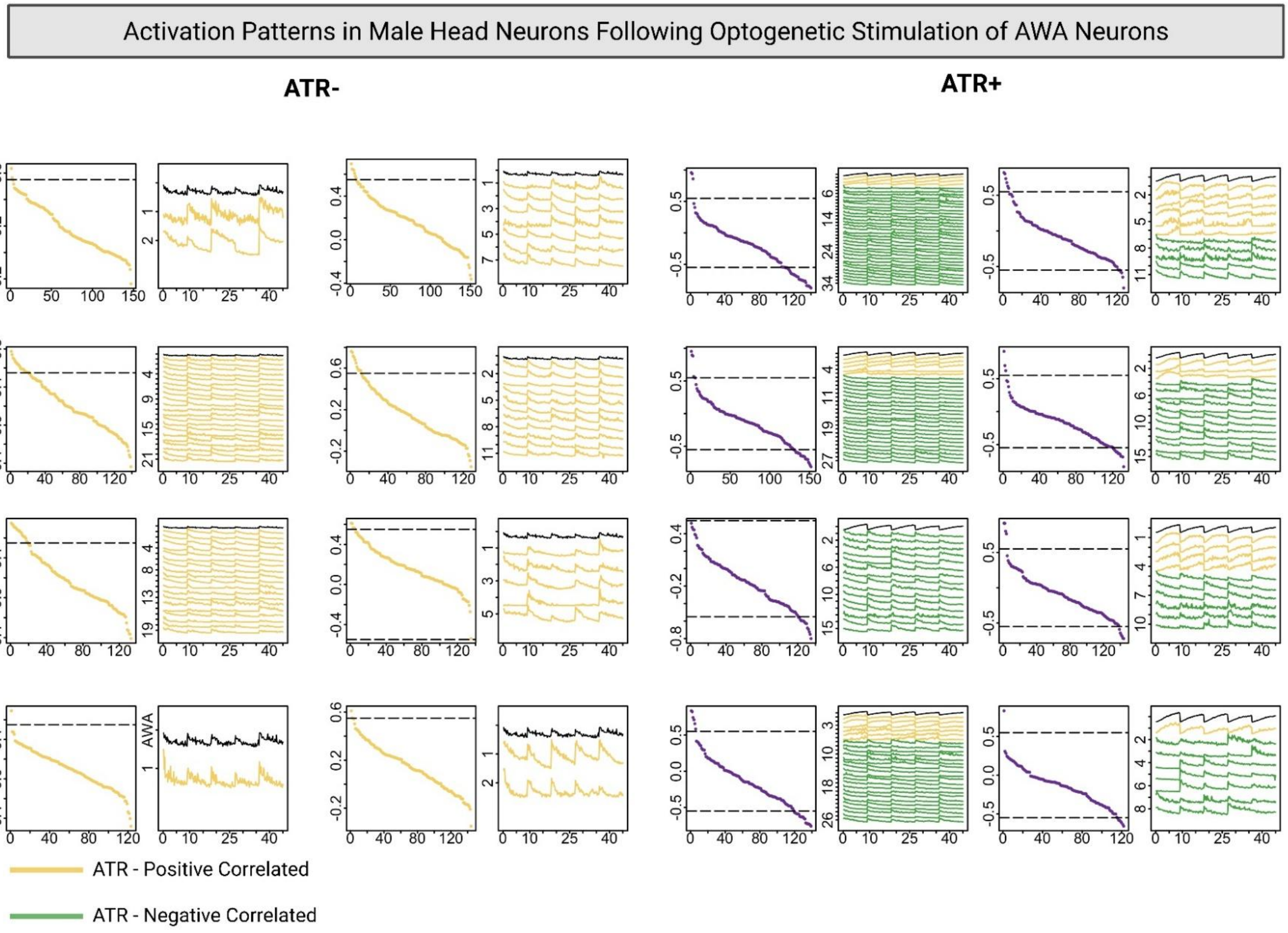


**Figure S9: Neural activity correlations with AWA and ASI optogenetic stimulation in head regions.** Yellow and purple indicate positive correlations with target neuron activity in ATR- and ATR+ samples, respectively. Green denotes negative correlations in ATR- and ATR+ samples, respectively. The dashed lines indicate the thresholds for positive and negative correlations. Sample sizes: control and experimental groups (n=8 each). Source data are provided as a Source Data file.

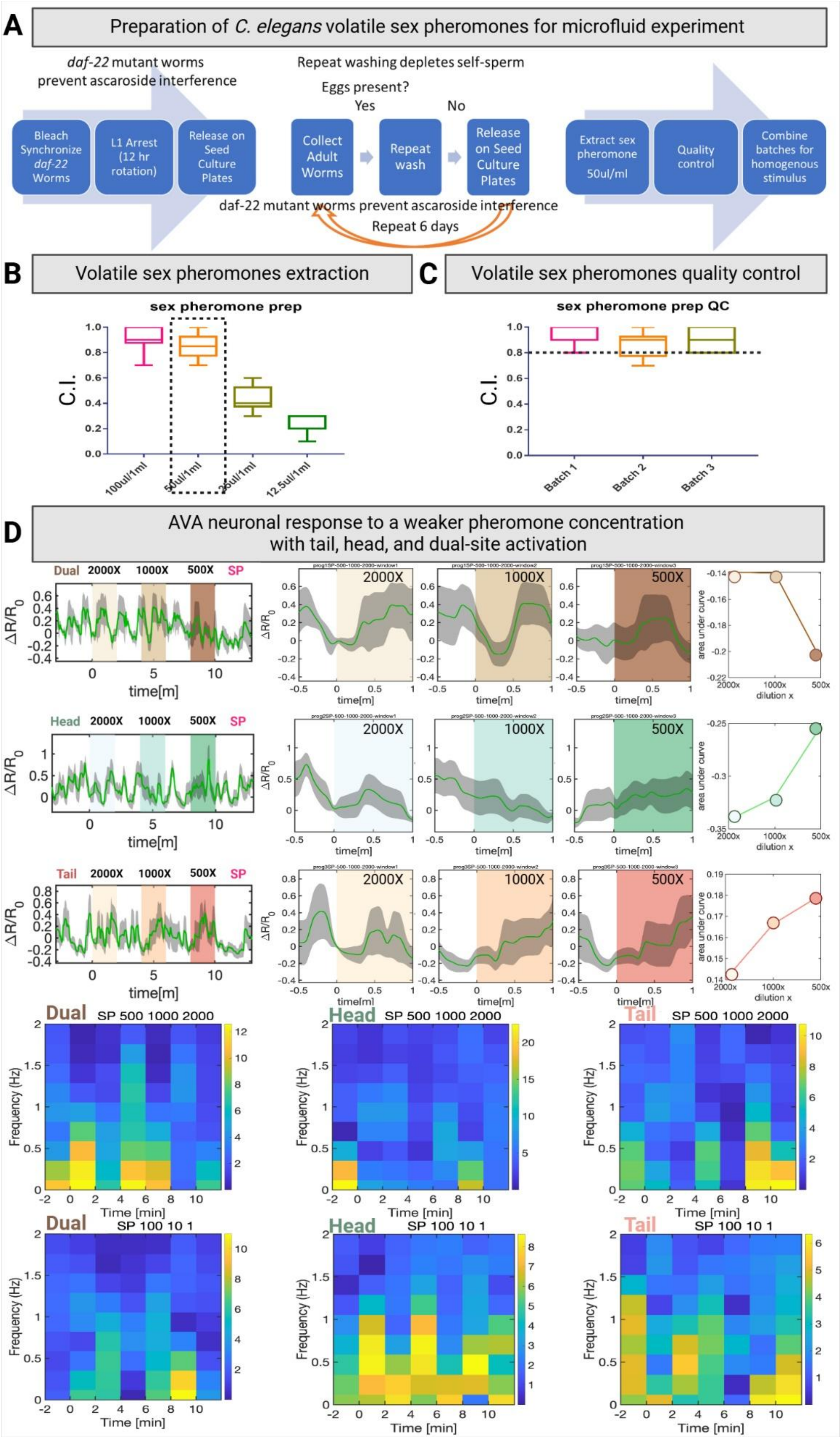
A
Preparation of C. elegans volatile sex pheromones for microfluid experiment
daf-22 mutant worms prevent ascaroside interference
Repeat washing depletes self-sperm
Eggs present?
Yes
No
Bleach Synchronize daf-22 Worms
L1 Arrest (12 hr rotation)
Release on Seed Culture Plates
Collect Adult Worms
Repeat wash
Release on Seed Culture Plates
Extract sex pheromone 50ul/ml
Quality control
Combine batches for homogenous stimulus
daf-22 mutant worms prevent ascaroside interference
Repeat 6 days
B
Volatile sex pheromones extraction
sex pheromone prep
C.I.
100ul/1ml
50ul/1ml
25ul/1ml
12.5ul/1ml
C
Volatile sex pheromones quality control
sex pheromone prep QC
C.I.
Batch 1
Batch 2
Batch 3
D
AVA neuronal response to a weaker pheromone concentration with tail, head, and dual-site activation
Dual 2000X 1000X 500X SP
Head 2000X 1000X 500X SP
Tail 2000X 1000X 500X SP
ΔR/R0
time[m]
area under curve
dilution x
Dual SP 500 1000 2000
Head SP 500 1000 2000
Tail SP 500 1000 2000
Dual SP 100 10 1
Head SP 100 10 1
Tail SP 100 10 1
Frequency (Hz)
Time [min]

**Figure S10: Sex pheromone preparation for microfluidics chip-based experiment. (A)** Modified Sex Pheromone Extraction Protocol for microfluidics chip-based experiments. To obtain large quantities of crude sex pheromone for microfluidics chip-based experiments, we streamlined the standard protocol. We used *daf-22* mutant *C. elegans* hermaphrodites, eliminating confounding ascaroside signals. Worms were synchronized and repeatedly washed to deplete self-sperm, and the sex pheromone extraction ratio was optimized. **(B)** Ratio Optimization: We tested four ratios (100µl/ml, 50µl/ml, 25µl/ml, 12.5µl/ml) for their attractiveness to males using a chemoattraction assay. The 50µl/ml ratio exhibited attractiveness comparable to 100µl/ml, with attractiveness declining at lower concentrations. Therefore, 50µl/ml was chosen for sex pheromone extraction. **(C)** Quality Control and Homogenization: Three independent batches of extracted pheromone were quality-controlled using the chemoattraction assay, all exceeding a predetermined quality threshold suggested by previous studies (C.I. = 0.8). Finally, the quality-controlled batches were combined to create a homogenous stimulus for microfluidics chip-based experiments. (B) and (C) Error bars represent the s.e.m. **(D)** Effects of sex pheromone stimulation on AVA reversal command interneuron activity in *C. elegans* males at Higher Dilutions. The calcium indicator GCaMP6s was used to monitor calcium dynamics in the AVA neuron. The AVA-specific split cGAL driver construct consisted of gpa-14P::NLS::cGAL (DBD)::gp41-1::N-intein::let-858 3'UTR and rig-3P::NLS::gp41-1::C-intein::cGAL(AD)::let-858 3'UTR, paired with the GCaMP effector construct 15xUAS::GCaMP7b::SL2::mKate2. The upper panel shows the calcium-shaded areas representing the s.e.m. This set of microfluidics chip-based experiments used 500X, 1000X, and 2000X fold diluted pheromones. The right panel shows area-under-curve (AUC) analysis (see Method). The bottom panel displays oscillation frequency distributions, revealing spontaneous oscillations in AVA neuron activity upon exposure to 500X, 1000X, and 2000X diluted pheromones. Heatmaps represent the frequency responses of AVA neurons under varying stimulation conditions. Each panel corresponds to specific activation sites (head, tail, and dual activation) and serial pheromone dilutions (set 1: 1X, 10X, 100X; set 2: 500X, 1000X, 2000X). The heatmaps depict the oscillation frequency dynamics of AVA activity over 12 mins for each stimulus program, with color coding indicating the distribution and intensity of oscillation frequencies. Each condition (head, tail, or dual location) at each concentration was tested with a sample size of 8 worms. Source data are provided as a Source Data file.

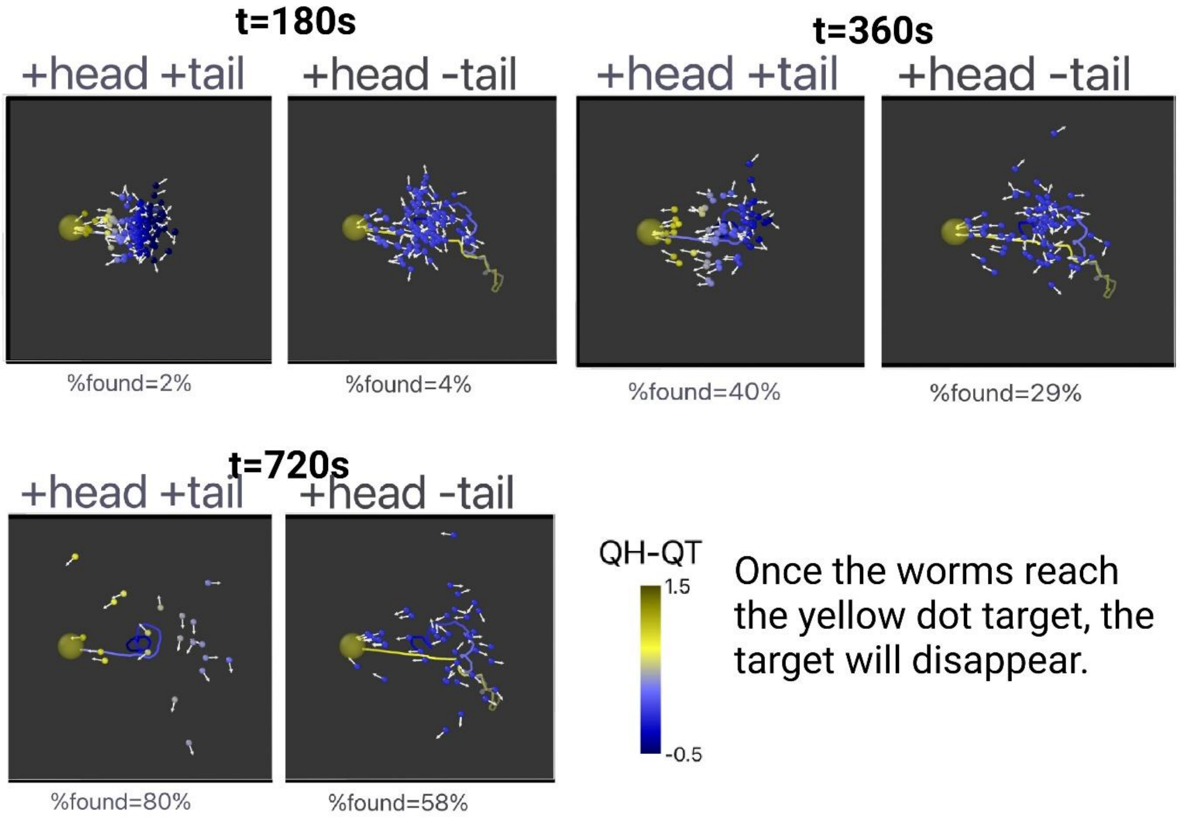
t=180s
+head +tail
+head -tail
%found=2%
%found=4%
t=360s
+head +tail
+head -tail
%found=40%
%found=29%
t=720s
+head +tail
+head -tail
%found=80%
%found=58%
QH-QT
1.5
-0.5
Once the worms reach the yellow dot target, the target will disappear.

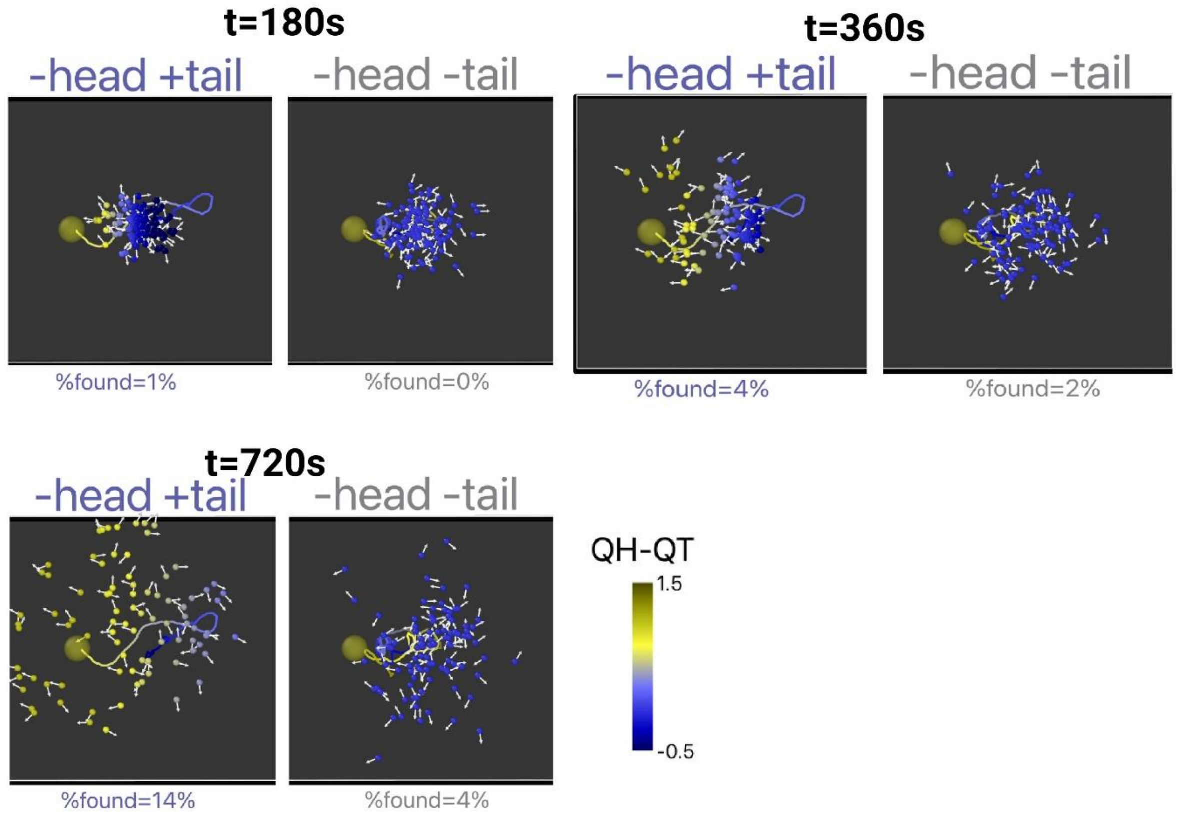
t=180s
-head +tail
-head -tail
%found=1%
%found=0%
t=360s
-head +tail
-head -tail
%found=4%
%found=2%
t=720s
-head +tail
-head -tail
%found=14%
%found=4%
QH-QT
1.5
-0.5

**Figure S11:** Each panel shows simulation snapshots comparing head/tail input-enabled conditions. In each snapshot with a time point, 100 randomly chosen worms from 10,000 stimulation events are displayed as spheres, colored by their instantaneous confidence $Q^H - Q^T$. Source data are provided as a Source Data file.

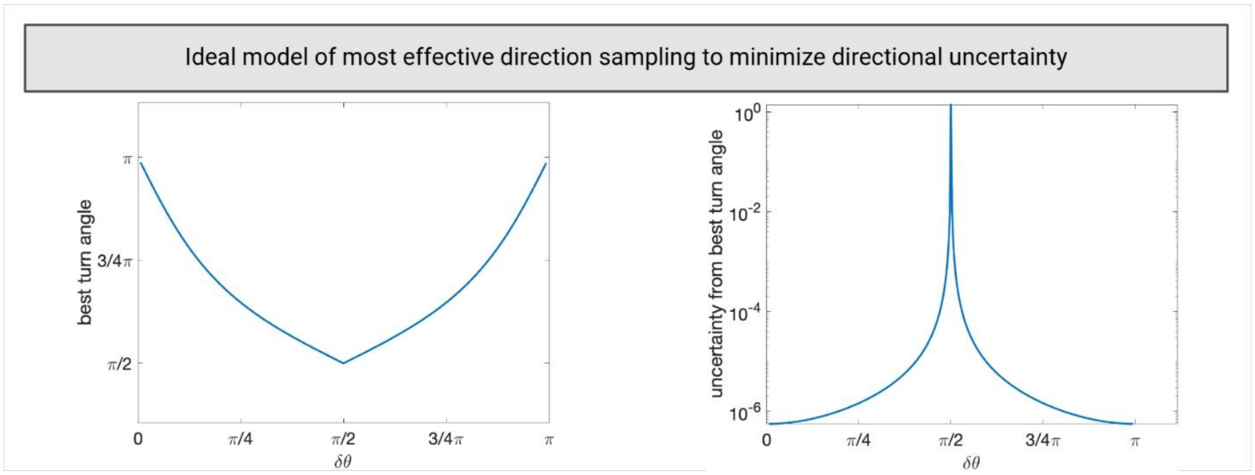


**Figure S12:** Ideal model for effective direction sampling to minimize directional uncertainty. δθ is the angle between the first sampled direction and the concentration gradient. The left panel shows the best angle to turn to sample the second direction. The right panel shows the directional uncertainty associated with the best turn angle. Averaging over an assumed uniform distribution of δθ results in a best turn angle close to 120 degrees.
